# A fully phased, chromosome-scale genome of sugar beet line FC309 enables the discovery of Fusarium Yellows resistance QTL

**DOI:** 10.1101/2024.05.04.592522

**Authors:** Olivia E. Todd, Sheron Simpson, Brian Scheffler, Kevin M. Dorn

## Abstract

Sugar beet (*Beta vulgaris* L.) is a global source for table sugar and animal fodder. Here we report a highly contiguous and haplotype phased genome assembly and annotation for sugar beet line FC309. Both assembled haplomes for FC309 represent the largest and most contiguous assembled beet genomes reported to date, as well as gene annotations sets that capture over 1500 additional protein-coding loci compared to prior beet genome annotations. These new genomic resources were used to identify novel quantitative trait loci (QTL) for Fusarium Yellows resistance from the FC309 genetic background using an F2 mapping-by-sequencing approach. The highest QTL signals were detected on Chromosome 3, spanning approximately 10 Mbp in both haplomes. A parallel transcriptome profiling experiment identified candidate genes within the Chromosome 3 QTL with plausible roles in disease response, including NBS-LRR genes with expression trends suggestive of their role as causal resistance genes. Collectively, the genomic resources for FC309 presented here be foundational tool for comparative genomics, mapping other traits in the FC309 background, and as a reference genome for other beet studies due to its contiguity, completeness, and high-quality gene annotations.

## 1. Introduction

The sugar beet (*Beta vulgaris* spp. *vulgaris* L.) genome has served as a powerful tool for global research and for breeding programs in both the public and private sectors (Dohm et al., 2014; McGrath et al., 2023). Both the RefBeet and EL10 assemblies have enabled the sugar beet research community to identify genetic loci linked to important traits underlying important agronomic, biotic, and abiotic stress tolerance traits (Dally et al., 2014; Galewski et al., 2022; Holmquist et al., 2021; Pfeiffer, 2016; Pfeiffer et al., 2017; Ravi et al., 2021a; Sielemann et al., 2023; Wigg et al., 2023). The rapid advancement in sequencing technologies has resulted in a moving target of benchmarks that represent ‘gold standard’ genome assembly and annotations for many species. The continued drastic reduction in sequencing costs, coupled with technologies like Hi-C (Belton et al., 2012; Putnam et al., 2016), and long read RNA sequencing make the assembly and annotation of small (<1Gb) diploid genomes an increasingly common exercise. Thus, the creation of new, purpose-built genomic resources to map traits from specific genetic backgrounds is easily within reach to avoid reference genome bias (Prasad et al., 2022; Thorburn et al., 2023). Additionally, widescale adoption of sequencing technologies that give both longer and more accurate reads has reliably improved contiguity and completeness (assembly size / estimated genome size) of assemblies. These more complete assemblies enable a deeper understanding of pan-genomic structural variation, characterization of repetitive element contraction, expansion, and genic conservation. The high degree of genome assembly completeness achieved with these new technologies changes the breadth of utility that ranges from identification of single genes of interest to comprehensive investigations of species-level population genetics (Thorburn et al., 2023).

Breeding targets in public sugar beet pre-breeding programs have historically focused on identifying resistance in sugar beet or beet crop wild relative germplasm (ex. *Beta vulgaris* subsp. *maritima,* or *Bvm* herein) (Panella et al., 2015). Compared to other crops, only recently has this focus shifted to molecular genetics and genomics-based research to identify markers associated with pathogen disease resistance traits (Ravi et al., 2021b; Stevanato et al., 2012; Tehseen et al., 2023). As genomic resources grow in sugar beet, there will likely be an increased effort to identify and extensively characterize resistance trait loci in existing sugar beet germplasm.

Among the resistance traits prioritized for sugar beet host plant fortification is resistance to the fungal pathogen *Fusarium oxysoprum.* This pathogen has been recognized as problematic for sugar beet in the United States (BSDF, 2021), and no effective chemical control options currently exist to manage this disease. Thus, improved host plant resistance represents the primary tool for growers to limit financial impact from this disease. The recent release of the sugar beet genetic stock FC309 (Todd et al., 2023), which has robust resistance to Fusarium Yellows derived from the *Bvm* background, represents a resurrected effort to genetically fix Fusarium resistance for breeders, genetically map, and characterize the resistance mechanism(s) at the molecular level.

In this study, we present a fully phased and annotated chromosome-scale assembly of sugar beet line FC309. We report assembly metrics and comparisons to other beet genome assemblies. This new genomic resource is used here to map and characterize a novel Quantitative Trait Loci (QTL) for Fusarium Yellows resistance using a mapping-by-sequencing, transcriptome profiling, variant effect analysis, and comparative genomics. Taken together, the results from these experiments suggest that nucleotide binding (NB) and leucine rich repeat (LRR) domain containing proteins known as NBS-LRRs contribute to Fusarium Yellows resistance in FC309.

## 2. Materials and Methods

### 2.1 FC309 Tissue Collection, Sequencing, Assembly, & Comparative Analyses

Seeds for USDA germplasm line FC309 (Todd et al., 2023) were planted and grown in 25°C, 16h day greenhouse conditions. Twelve individuals in differing growth stages were collected to maximize transcript discovery for RNA isolation for PacBio IsoSeq. Untreated cotyledon and two-leaf stage seedlings were flash frozen in liquid nitrogen. Six eight-leaf stage plants were separated into root and shoot tissue types and flash frozen 30 days after inoculation with *Fusarium oxysporum* strain F19 as described in Todd et al. (2023). Six additional *Fusarium* treated eight-leaf stage individuals were placed in a 4C room to vernalize for 190 days following the 30 days after inoculation period, separated by roots and shoots and flash frozen in liquid nitrogen 14 days after being removed from vernalization. RNA was extracted from individual plants using a QIAGEN RNeasy extraction kit. Equimolar pools of RNA from each of the above-mentioned tissue types were combined to create one sample for IsoSeq library preparation and sequencing at the University of Minnesota Genomics Center (MN, USA) on the PacBio Sequel II platform.

For genome sequencing, one treated FC309 individual was taken from the aforementioned planting and placed into a light-impermeable cabinet for three days. Approximately 3g of young, dark-treated tissue was harvested into 50mL falcon tubes and flash frozen in liquid nitrogen. A PacBio Circulomics high molecular weight DNA extraction kit was used to obtain DNA samples for PacBio HiFi sequencing. Three total SMRT cells (PacBio Sequel II platform) worth of HiFi sequencing were produced from this FC309 individual plant. Over 64 Gb of total HiFi data were generated for this single FC309 individual, or >86X genome coverage.

DoveTail Omni-C reads were obtained by submitting 3g of fresh, dark treated tissue to DoveTail Genomics for Omni-C library creation and Illumina sequencing. The tissue was taken from the same individual that the HiFi sequencing was obtained from. Chromatin was fixed in place with formaldehyde in the nucleus and then extracted. Fixed chromatin was digested with DNase I, chromatin ends were repaired and ligated to a biotinylated bridge adapter followed by proximity ligation of adapter containing ends. After proximity ligation, crosslinks were reversed, and the DNA purified. Purified DNA was treated to remove biotin that was not internal to ligated fragments. The sequencing library were generated using NEBNext Ultra enzymes and Illumina-compatible adapters. Biotin-containing fragments were isolated using streptavidin beads before PCR enrichment of the library. The library was sequenced on an Illumina HiSeqX platform (2×150 bp reads) to produce 18.9 Gb of total data.

The HiFi reads and Dovetail Omni-C reads were used as input data for phased assembly with hifiasm (v0.19.4, Cheng et al., 2021). Omni-C reads were quality filtered and trimmed to remove adaptor contamination using BBduk (v39.01, part of the BBTools package, sourceforge.net/projects/bbmap/) using the following settings: k=23, ktrim=r, qtrim=rl, trimq=20, mink=11. HiFi reads were trimmed and filtered to remove adaptor contamination using HiFiAdapterFilt (v2.0.1, Sim et al., 2022). The resulting phased contig-level assemblies were subsequently analyzed with the NCBI tools fcs-adaptor and fcs-gx to identify sequencing adaptor and foreign DNA contamination. These analyses identified no contamination, and the contig sets from each phased assembly were used as input for HiRise scaffolding (Putnam et al, 2016). HiRise is a computational pipeline designed specifically for using proximity ligation data to scaffold genome assemblies. Briefly, filtered Dovetail OmniC library sequences were aligned to the draft input assembly using bwa (https://github.com/lh3/bwa). The phased haplomes from the Dovetail Omni-C read pairs mapped within draft scaffolds were analyzed by HiRise to produce a likelihood model for genomic distance between read pairs. The likelihood model was used to identify and break putative mis-joins, score prospective joins, make joins above a threshold, and assemble nine major scaffolds. Unanchored scaffolds (i.e., those not assigned to a pseudo-chromosome) were concatenated with the 9 phased psuedochrosomomes in each respective phased assembly. The haplome 1 assembly was named “USDA_Bvulg_FC309_v1.1.0” (NCBI accession JAYERX000000000) and the haplome 2 assembly was named “USDA_Bvulg_FC309_v1.2.0” (NCBI accession JAYERY000000000).

Pseudo-chromosomes (9 largest scaffolds) from each FC309 haplome were first reoriented to match the linear orientation of the RefBeet v1.2.2 psuedo-chromsomes to enable synteny visualization. The FC709-3 v1.0.1 pseudochromosomes were aligned to both the RefBeet and re-oriented EL10.2 pseudochromosomes using minimap2 (v2.24, Li 2018). The resulting BAM alignment files were indexed using the samtools index command (v1.16.1). Comparisons of the resulting alignments were processed using Syri (https://github.com/schneebergerlab/syri, version 1.6.3, Goel et al., 2019), implemented using Conda (v4.13.0). Visualization of alignments was implemented using plotsr (https://github.com/schneebergerlab/plotsr, v0.5.4, Goel and Schneeberger, 2022). Synteny and duplication visualizations presented in the Circos diagram (Figure 1) are from Syri.

**Figure 1:**
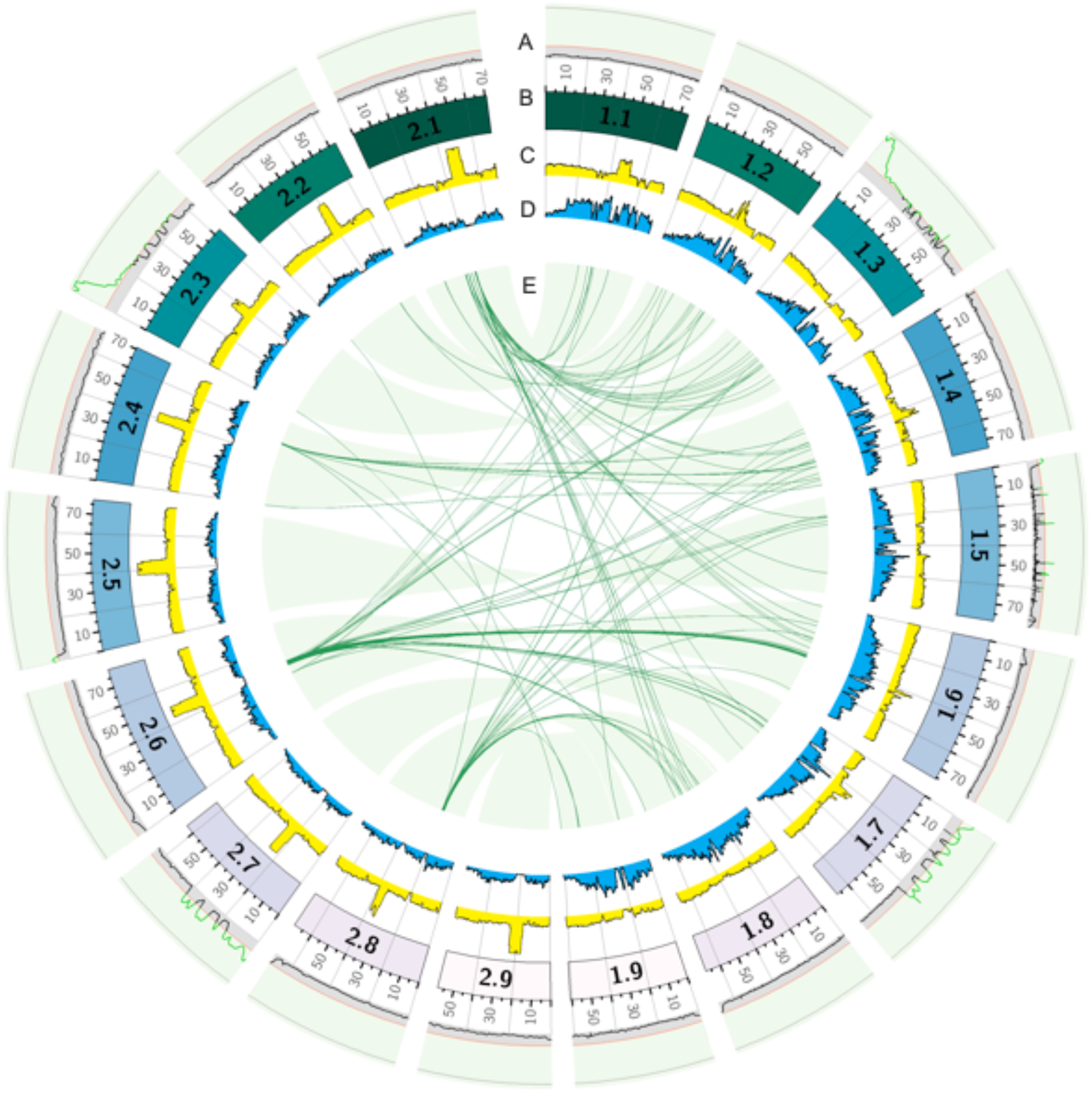
Circos graph showing (A) G’ value as determine by QTLseqR, where black line regions represent non-significant regions and bright green lined regions are statistically significant QTL, per chromosome. The cutoff for significant QTL is at y=8 (red line) and the top grey line is y = 38. (B) Karyotype for each n=9 haplome, where chromosome notation is x.y (x=haplome number, y=chromosome number), major tick marks are every 5 Mbp and the extended grid is represented in 20 Mbp increments. (C-D) Histograms of LTR (C) and unclassified (D) repeat distributions in a 0.5Mbp sliding window. (E) syntenic regions between n=9 chromosomes prefixed with either haplome 1 or 2. Light green ribbons show syntenic regions, dark green ribbons show inverted and non-inverted duplication locations.

### 2.2 FC309 Haplome Annotation

Removal of polyA tails and artificial concatemers in the PacBio IsoSeq reads was conducted with isoseq3 refine (v6.0.0). The resultant bam files for FC309 were aligned to the respective indexed haplomes using pbmm2 (v1.13.0). Redundant transcripts after aligning were collapsed with isoseq3 collapse (v6.0.0). The aligned, collapsed IsoSeq reads were uploaded to GenSAS and used with BRAKER2. Haplomes derived from sequencing were structurally and functionally annotated used GenSAS (Humann et al., 2019). Repeat databases for both haplomes of FC309 and FC709-3 were made using RepeatModeler (Flynn et al., 2020). RepeatMasker was set to mask interspersed and simple repeats using the rmblast engine. BRAKER2 (Hoff et al., 2019) was run in GenSAS with the HiFi IsoSeq transcript reads aligned to each haplome as evidence. Consensus gene sets were made from blastn alignments with both of the publicly available EL10.2_2 transcript data and Refbeet-1.2.2 genomic sequences (Dohm et al., 2014; McGrath et al., 2023) and the BRAKER2 output using EvidenceModeler (Haas et al., 2008). The official gene set (OGS) was refined using PASA (Haas et al., 2011). Functional annotation was completed on the refined dataset with InterProScan (v5.53-87.0) and Pfam (version 1.6).

### 2.3 Transcriptome Profiling with RNA-sequencing

Seeds for USDA germplasm line FC309 (Todd et al., 2023) and USDA germplasm line Monohikori (susceptible to *Fusarium oxysporum* strain F19) were planted in steam pasteurized soil with 4 inch pots and grown in 25C, 16h day greenhouse conditions. Four individuals per line were sampled at an untreated timepoint, 24 hours post-inoculation and 6 days post-inoculation. Plants were treated via the root dip method described by Todd et al. (2023). The untreated plants were dipped in a solution of carboxymethylcellulose (CMC) for 5 minutes with no *Fusarium oxysporum* strain F19 spores present. All plants upon harvest were flash frozen in liquid nitrogen and RNA was extracted using a QIAGEN RNeasy plant minikit and the samples were treated with TURBO DNase (Thermo Fisher Scientific) prior to sample submission. All samples passed initial QC for Illumina HiSeq4000 except for two: individuals one and two from the untreated FC309 timepoint. These samples were included in the sequencing. The .fastq files were aligned to the indexed FC309 haplomes using hisat2 (v2.1.0) (Zhang et al., 2021) (SI Table 2). Counts were tabulated using FeatureCounts from the Subread package (v2.0.6) (Liao et al., 2013). The counts file was analyzed using Deseq2 in R (v1.36.0) (Love et al., 2014) to obtain differential expression profiles for a number of treatment comparisons.

### 2.4 Variant Calling, QTL and Synteny Analysis

Biparental mapping was conducted by crossing the FC309 resistant line with FC308, a USDA released germplasm that is highly susceptible to *Fusarium oxysporum* strain F19 (Todd et al., 2023). One F2 line was treated with *Fusarium oxysporum* strain F19 in the same manner described in the RNA sequencing experiment (n = 123). Three phenotypic classifications (per Todd et al, 2024) were assigned to each plant following treatment: highly resistant (<29% symptom injury or less), intermediate (30-79%) and highly susceptible (>80% injury). Pools of 50 highly resistant and 50 highly susceptible DNA were sequenced with Illumina short read technology. Resistant and susceptible pools were mapped to each of the FC309 haplomes using Hisat2 (v2.2.1). The resulting SAM format mapping files for each pool converted, sorted, and indexed using SamTools (v1.17). Variant calling was conducted using BCFtools (v1.16) for both the resistant and susceptible pools for each haplome. A final variant table was created using the GATK tool VariantsToTable, which was used as input for the QTLseqr package (v.0.7.5.2, Mansfeld and Grumet, 2018) in R (v3.6.1, R Core Team, 2019). Within QTLseqR, the ‘Fusarium Susceptible’ bulk was assigned to the ‘highBulk’ variable and the ‘Fusarium Resistant’ bulk sample was assigned to the ‘lowBulk’ variable using the importFromGATK function. The following settings were used for each of the stated functions within QTLseqR: filterSNPs: refAlleleFreq = 0.2, minTotalDepth = 50, maxTotalDepth = 1000, minSampleDepth = 25, minGQ = 99, runGprimeAnalysis: windowSize = 1e6, outlierFilter = “deltaSNP”, filterThreshold = 0.2, and runQTLseqAnalysis: SNPset = df_filt, windowSize = 1e6, popStruc = “F2”, bulkSize = c(50, 50), replications = 10000, intervals = c(95,99). QTL peaks were chosen for analysis based on highest tricube smoothed G statistic (G Prime) (SI Figure 1). The delta SNP index value, the difference in proportion of alternate alle depth and total read depth per bulk was also calculated with 95% and 99% confidence intervals (SI Figure 1).

## 3. Results and Discussion

### 3.1 Genome Assembly, Annotation, and Comparative Genomics

The genome size was found to be approximately 633 Mbp for haplome 1 and 619 Mpb for haplome 2, not including unanchored contigs (Table 1). The estimated scaffold N/L50 was 5/67.76 Mbp for haplome 1 and 5/69.743 Mbp for haplome 2 (Table 1). Out of 30,170 protein coding genes annotated on the 9 chromosomes for each haplome, there were 913 protein coding genes on the unanchored contigs of haplome 1 and 29,551 protein coding genes on 9 chromosomes of haplome 2 (213 protein coding genes on the unanchored contigs of haplome 2) (Table 1). The assembled haplome size is approximately 9% larger than Refbeet, a doubled haploid reference genome (Dohm et al., 2014) and ∼13.2% larger than EL10.2_2, a recent reference constructed with PacBio sequel II long read technology, Hi-C scaffolding and polished with Illumina paired-end reads (McGrath et al., 2023). Individual chromosome sizes ranged from 60-78 Mbp for haplome 1 and 58-79 Mbp for haplome 2 (Figure 1B, Table 1). BUSCO scores for the generated haplomes were 94.3% and 89%, respectively. We generated 64.6, 18.9 (126M reads) and 7.7 (4.3M reads) gigabases of sequencing data on FC309 for HiFi, Omni-C and IsoSeq, respectively.

**Table 1.**
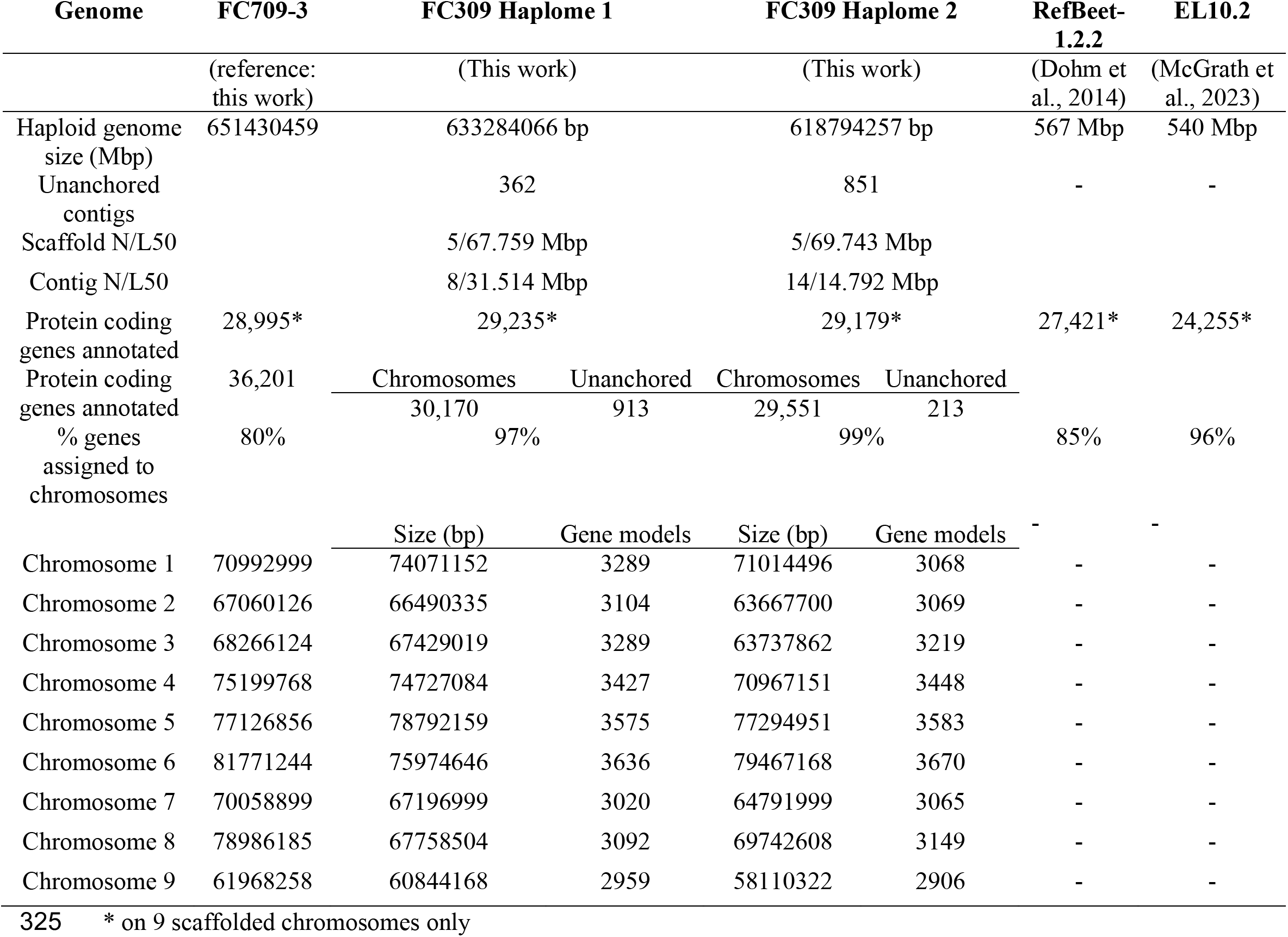
Comparison of genomic features and assembly statistics between the most recent two genome releases and currently available sugar beet genomes.

We predicted that approximately 66% of each haplome was covered by repetitive sequences. The largest portion of the genome was covered by unclassified repeat elements (40.04% and 38.44% for haplome 1 and 2 respectively) (Supplemental Table 1). The unclassified repeats were generally uniformly distributed over all 9 chromosomes, but showed an increase near the centromere in both haplomes (Figure 1C-D). Long terminal repeat (LTR) retrotransposons covered 103 Mbp (15.87%) and 105 Mpb (15.78%) of haplome 1 and haplome 2, respectively (Supplementary Table 1), and were found to be less present near the telomeres of all 9 chromosomes for both haplomes (Figure 1C). The LTR elements were the largest repeat element group following the unclassified group, mainly comprised of Ty1/Copia and Gypsy/DIRS1 repeats. This report contrasts the results from EL10.2_2, where the largest repeat element group was reported to be DNA transposons (58%), but support the findings reported in RefBeet (Dohm et al., 2014; McGrath et al., 2023). Overall, the proportion of repeat elements reported is congruent with previous studies in beet and related crops (Flavell et al., 1974) and sequence coverage analysis showed high coverage over areas of estimated high repeat density, a benefit of long-read sequencing (Putnam et al., 2016).

Whole-genome alignments between the two FC309 haplomes identified each assembly were largely syntenic (Figure 1, track E, light green ribbons). However, notable inversions (Chromosome 1), presence/absence variation (Chromosomes 3, 4, 8, and 9), and duplications (Chromosome 6) were identified (Supplemental Figure 1). Alignments of the FC309 haplomes to RefBeet and EL10 identified substantial genomic rearrangements and duplications (Supplemental Figure 1), highlighting the high likelihood of reference genome bias when using these reference genomes for mapping traits of interest in the FC309 background.

### 3.2 Quantitative Trait Loci (QTL) Identification, Transcriptome Profiling, and Mapping-by-Sequencing

A total of 19,305 and 18,824 SNPs were detected for haplome 1 and haplome 2, respectively in the top QTL region identified with QTLseqR. This major effect QTL on chromosome 3 that spanned 30 Mbp (Figure 1A) and had a max G’ of ∼36 for both haplomes, which was approximately 2 times larger than the next highest peaks on chromosome 7 with a G’ values of ∼17.8. Other minor effect QTL were found on chromosome 7 of haplome 1 and 2 (Supplemental Figure 2). These QTL were identified by the tri-cubed ΔSNP index and smoothed G’ statistics, which have been shown to improve accuracy of causative SNP detection by as much as 44% for species with low-recombination rates compared to the non-smoothed statistic (de la Fuente Cantó and Vigouroux, 2022). The peak delta SNP position at the chromosome 3 QTL region was 3,686,961 bp for haplome 1, and 3,985,697 bp for haplome 2.

We mapped expression trends and characterized candidate genes approximately ± 1 Mbp of these genomic coordinates, spanning position 2,406,020 to 4,601,366 containing 182 genes (haplome 1) and 2,923,159 – 4,940,399 containing 175 genes (haplome 2) (Figure 2). Seven disease resistance proteins were present in the QTL region for both haplomes, which were the primary candidates to explore transcript expression and SNP investigation. Six of these proteins are spatially oriented after one another, with Bv.00g064600.m01 and Bv.00g064610.m01 in one cluster with opposite orientations, and Bv.00g065610.m01, Bv.00g065630.m01, Bv.00g065640.m01 and Bv.00g065650.m01 in another cluster, all with reverse strand orientation (haplome 1 nomenclature). Interestingly, while it may be possible that all four disease resistant proteins in cluster 2 are under the same promoter due to proximity, their expression remains unchanged in response to treatment with *Fusarium*, with the exception of Bv.00g065650.m01/Bv.00g065330.m01 (haplome 1/haplome 2), which shows constitutive expression at the untreated treatment in FC309, but has extremely low transcript expression in the susceptible line (Figure 3). The singular disease resistance gene Bv.00g064370.m01/Bv.00g063890.m01 (haplome 1/haplome 2) was constitutively expressed in FC309 and significantly upregulated at 24 HPI in FC309 compared to the susceptible. Interestingly, the transcript response begins to statistically increase in the susceptible line 6 days post inoculation (DPI) compared to the susceptible untreated condition (Figure 4A), suggesting a delayed transcription of a potentially necessary gene contributing to the resistance response.

**Figure 2:**
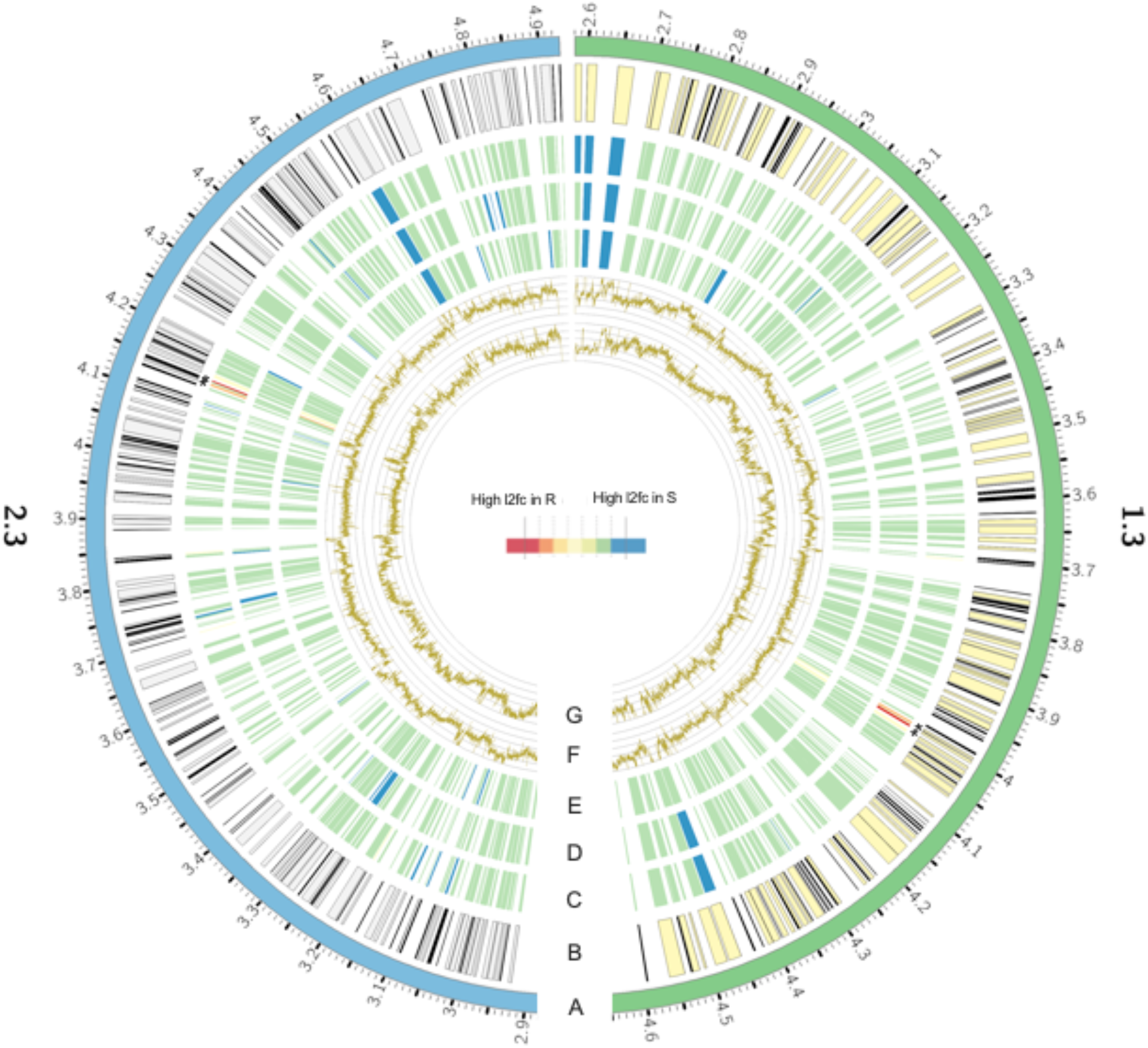
2 Mbp QTL regions on Chromosome 3 for haplome 1 (1.3) and haplome 2 (2.3). (A) Karyotype information for the 2 Mbp QTL region of haplome 1 and 2. (B) Coding sequence regions where coding regions greater than 3kb are represented by yellow and grey boxes (haplome 1 and 2, respectively) and black boxes for coding regions less than 3kb (both haplomes). (C-E) Heatmap of log2 fold change for three comparisons of interest: (C) The untreated condition for the resistant line versus the susceptible line. (D) The resistant line at 24 hours after treatment (HAT) compared to the susceptible line 24HAT and (E) the resistant line 6 days post inoculation (DPI) compared to the susceptible line 6DPI. Blue coloration equates to statistically significant upregulated transcript expression in the susceptible line, green indicates non-significant differential expression, and red indicates statistically significant upregulated transcript expression in the resistant line, per comparison. Asterisks (*) mark two candidate resistant genes. (F-G) Read coverage for Illumina sequencing of the F2 resistant pool (F) and the susceptible pool (G) for both haplomes. Grey Y intervals depicted correspond to 100 bp sliding windows, Ymax = 500X coverage.

**Figure 3:**
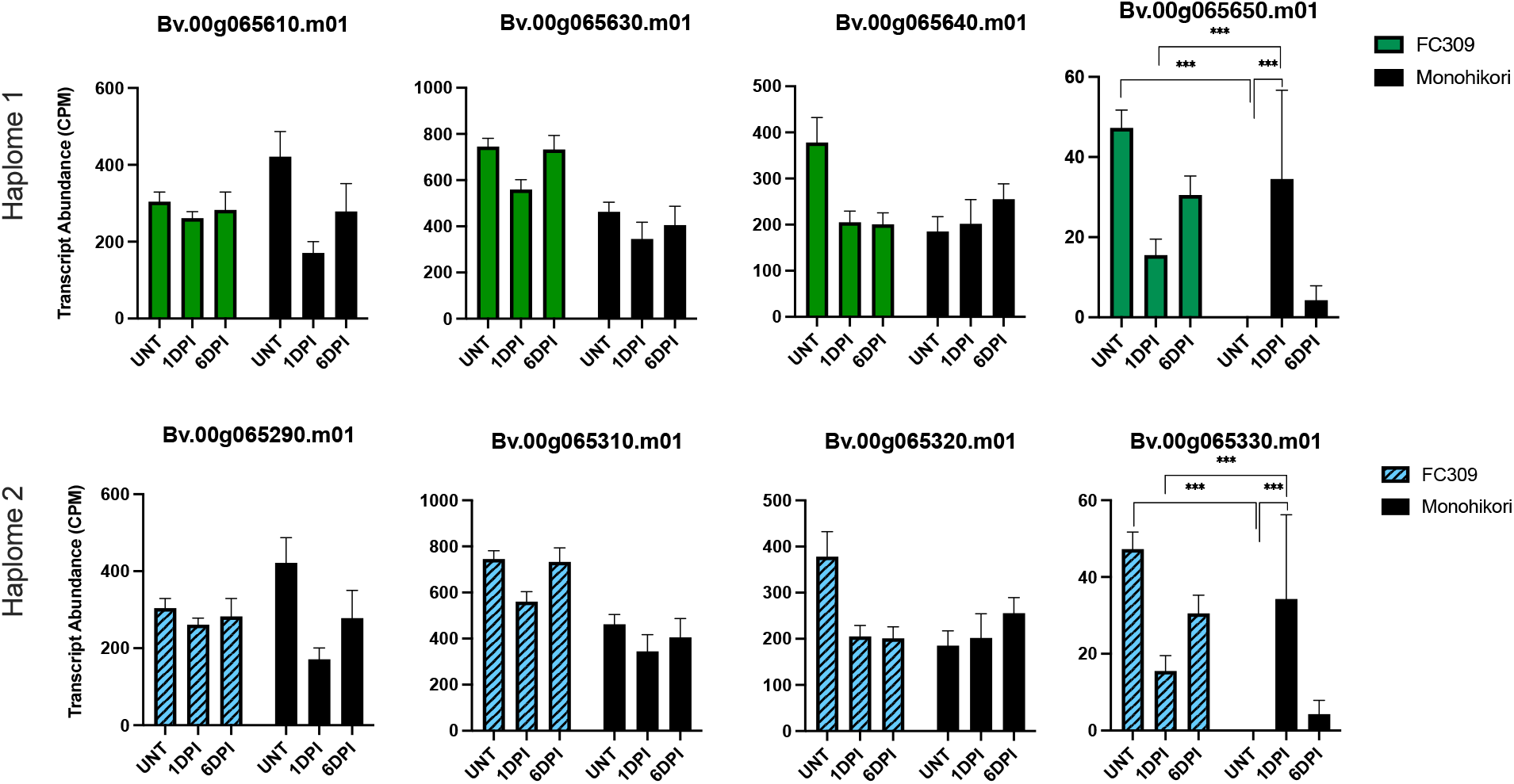
Candidate homologue transcript expression profiles for genes of interest in NBS-LRR domain containing proteins in cluster three of the haplome 1 (top graph panels) and haplome 2 (bottom graph panels) QTL intervals. Transcript expression is expressed as counts per million (CPM). Statistically significant comparisons are shown by asterisks at p= <0.001. For clarity, nonsignificant comparisons are not shown. The 24 hours post inoculation treatment is shown as 1 day post inoculation (1 DPI).

**Figure 4:**
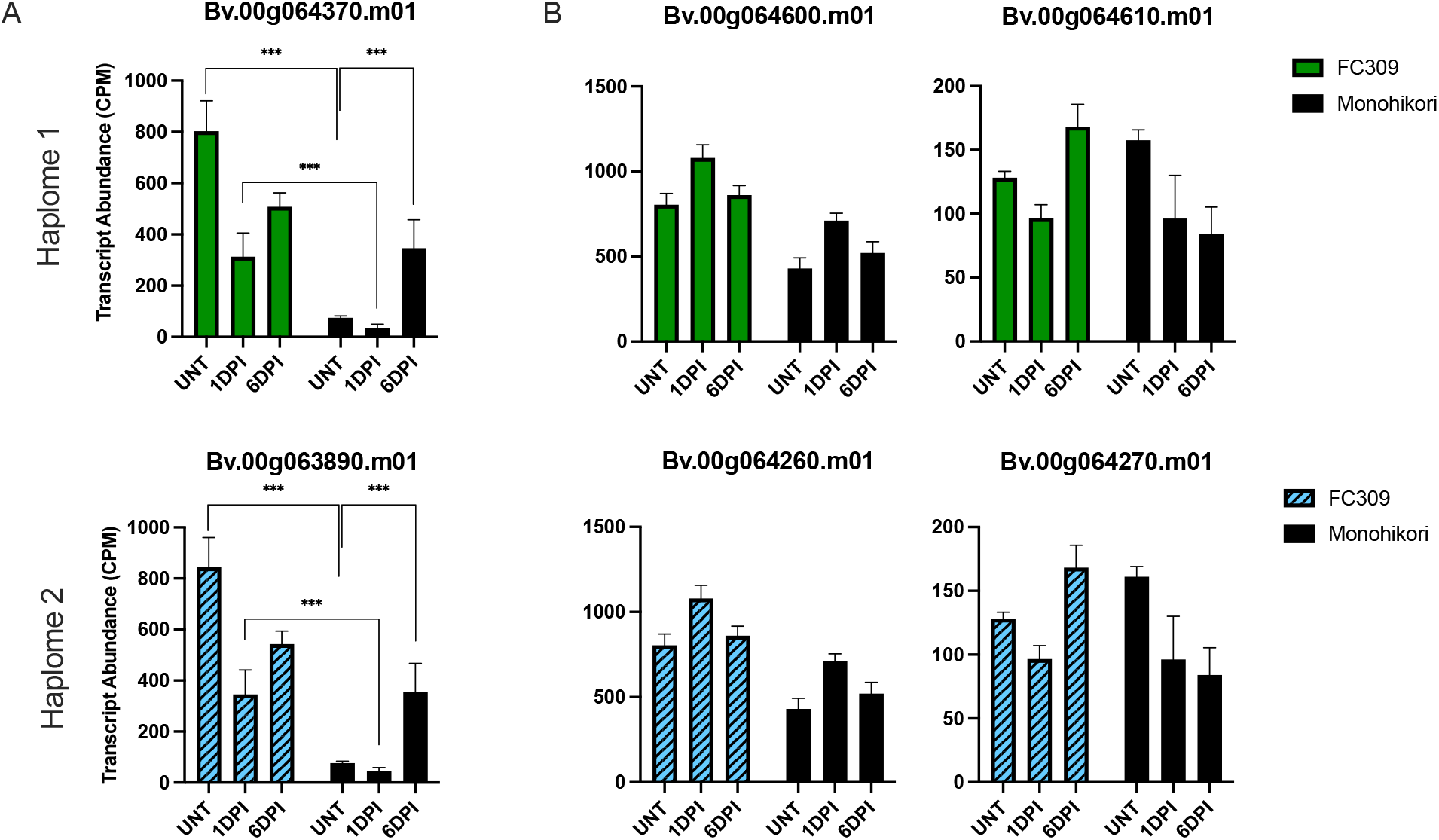
Candidate homologue transcript expression profiles for genes of interest in NBS-LRR domain containing proteins for (A) the singular gene and (B) cluster two of the haplome 1 (top graph panels) and haplome 2 (bottom graph panels) QTL intervals. Transcript expression is expressed as counts per million (CPM). Statistically significant comparisons are shown by asterisks at p= <0.001. For clarity, nonsignificant comparisons are not shown. The 24 hours post inoculation treatment is shown as 1 day post inoculation (1 DPI).

Both Bv.00g065650.m01/Bv.00g065330.m01 and Bv.00g064370.m01/Bv.00g063890.m01 encode NBS-LRR domain containing proteins (NBS-LRRs). NBS-LRRs function to identify pathogen effector proteins or detect pathogen associated proteins (DeYoung and Innes, 2006). NBS-LRRs have been implicated in *Fusarium oxysporum* resistance in various crops (Cao et al., 2024; Chakraborty et al., 2018; Liu et al., 2019), but have yet to be empirically related to *Fusarium* resistance in sugar beet. The Bv.00g065650.m01/Bv.00g065330.m01 homologues were not annotated as a specifically characterized disease resistance protein but was 89% homologous to RGA4 from *Chenopodium quinoa* from BLASTP results. Bv.00g065650.m01/Bv.00g065330.m01 contains CC, NB-ARC and LRR domains, which are a characteristic of the CNL class of NBS-LRR protein (Belkhadir et al., 2004; McHale et al., 2006). The Bv.00g065650.m01/Bv.00g065330.m01 homologues had no identified upstream, downstream or intergenic variants. Bv.00g064370.m01/Bv.00g063890.m01 encodes RPP8, a protein that has had a long-described role as a disease resistance protein in the gene-for-gene resistance model against oomycetes, but little is reported regarding the role of RPP8 in resistance to filamentous fungi (Cooley et al., 2000). RPP8 has been thought to not involve any other co-factors during the resistance response, somewhat atypical to the “guardee” hypothesis ((McDowell et al., 2000) reviewed by Glazebrook (2001)). Mutations in the W box *cis* elements of the promoter have been shown to affect *RPP8* expression in Arabidopsis, however the reported mutations caused reduced expression, not constitutive expression as we show in Bv.00g064370.m01/Bv.00g063890.m01 (Figure 4A) (Mohr et al., 2010). The Bv.00g064370.m01/Bv.00g063890.m01 homologues contain multiple sequence variants with two to eight “high” effect variants annotated by SnpEff. Numerous other moderate effect variations in upstream, downstream and intergenic regions were also detected that may also affect transcript expression or protein function. Exhaustive analysis and functional validation of the detected variants in RPP8, or other candidate genes identified here, will be pursued in future studies. While both QTL mapping and expression profiling results implicate this RPP8 as putatively causative, functional validation studies are needed to definitively ascertain the effects of the allelic variation identified here.

Numerous studies describe markers linked to *Fusarium oxysporum* resistance in various agronomic crops (Anjani et al., 2018; Deol et al., 2022; Jaber et al., 2020; Jha et al., 2021), and specifically NBS-LRRs (Cao et al., 2024; Chakraborty et al., 2018; Liu et al., 2019). De Lucchi et al. (2017) identified SNP markers in resistance gene analogs, including NBS-LRRs, linked to Fusarium resistance in sugar beet germplasm. This study did also utilize a progenitor line to FC309, however, the variants identified in this study were on chromosomes 2 and 7. This finding overlaps with minor QTL peaks identified here on chromosome 7 in both haplomes of FC309.

Analyses like co-expression network or pathway enrichment using the transcriptome profiling datasets describe here are outside the scope of this study but would provide useful new information to develop a deeper understanding of molecular genetic interactions between *Fusarium* and sugar beet. For example, in addition to NBS-LRRs, the MAP kinase pathway and WRKY transcription factors have also been implicated in resistance *Fusarium oxysporum* in other crops (Diao et al., 2021; Wang et al., 2018). However, the potential role of WRKY genes and the MAPK pathway has yet to be investigated in beet. Additional candidate genes and resistance mechanism pathways would likely be identified with these analyses.

## Conclusion

The ‘purpose-built’ FC309 sugar beet genome has enabled the direct discovery of a novel QTL linked to Fusarium Yellows resistance, and directly identified candidate genes and potentially causative variants from this background, notably *RPP8* and *RGA4*. Further experiments are needed to prove causation of any specific variant in these, or other, candidate genes. However, the rapid and precise identification of this QTL, and the underlying genes in the QTL interval, described here represent a substantial shift in genomics-assisted trait identification in sugar beet. Additional fine mapping experiments are ongoing using additional mapping populations to refine the QTL identified here. Future studies are needed to functionally validate the putative role of *RPP8, RGA4*, and other candidate genes in this QTL interval via transgenesis, protein modeling, pathogen-interaction experiments. As research in sugar beet, and other less-studied crops, continues shifting towards genomics-enabled characterization of specific plant-pathogen interactions, the results of this study demonstrate how rapidly specific traits can be mapped and candidate genes identified. Collectively, these results represent a first step in an emerging pipeline to comprehensively map disease resistance traits already present in beet breeding lines, with the long-term goal of exhaustively identifying novel traits present in wild beet germplasm.

## Data availability

The genomic resources described in this paper, including all raw sequencing reads, have been deposited at NCBI. Individual NCBI BioSample accessions are listed in Supplemental Table XX. Raw sequencing reads (PacBio HiFi and DoveTail Omni-C) for the FC309 genome assembly are available under BioProject accessions PRJNA1051123 (v1.1.0) & PRJNA1051124 (v1.2.0). Illumina reads from the F2 phenotypic bulks, Illumina transcriptome profiling, and PacBio IsoSeq are all available under PRJNA1051123. The phased assemblies for the FC309 genome are available at NCBI under accession numbers JAYERX000000000 (v1.1.0) and JAYERY000000000 (v1.2.0).

## Acknowledgements

This work was supported by USDA-ARS Appropriated Project Number 3012-21220-011-000D. The Western Sugar Joint Research Committee contributed partial funding for this project. The USDA is an equal opportunity provider and employer. Mention of trade names or commercial products in this publication does not imply recommendation or endorsement by the USDA.

## Author Contributions

OET Investigation, writing original draft, review and editing; SS and BS Investigation, review and editing; KMD Investigation, writing original draft, review and editing.

---

**Supplemental Figure 1:**
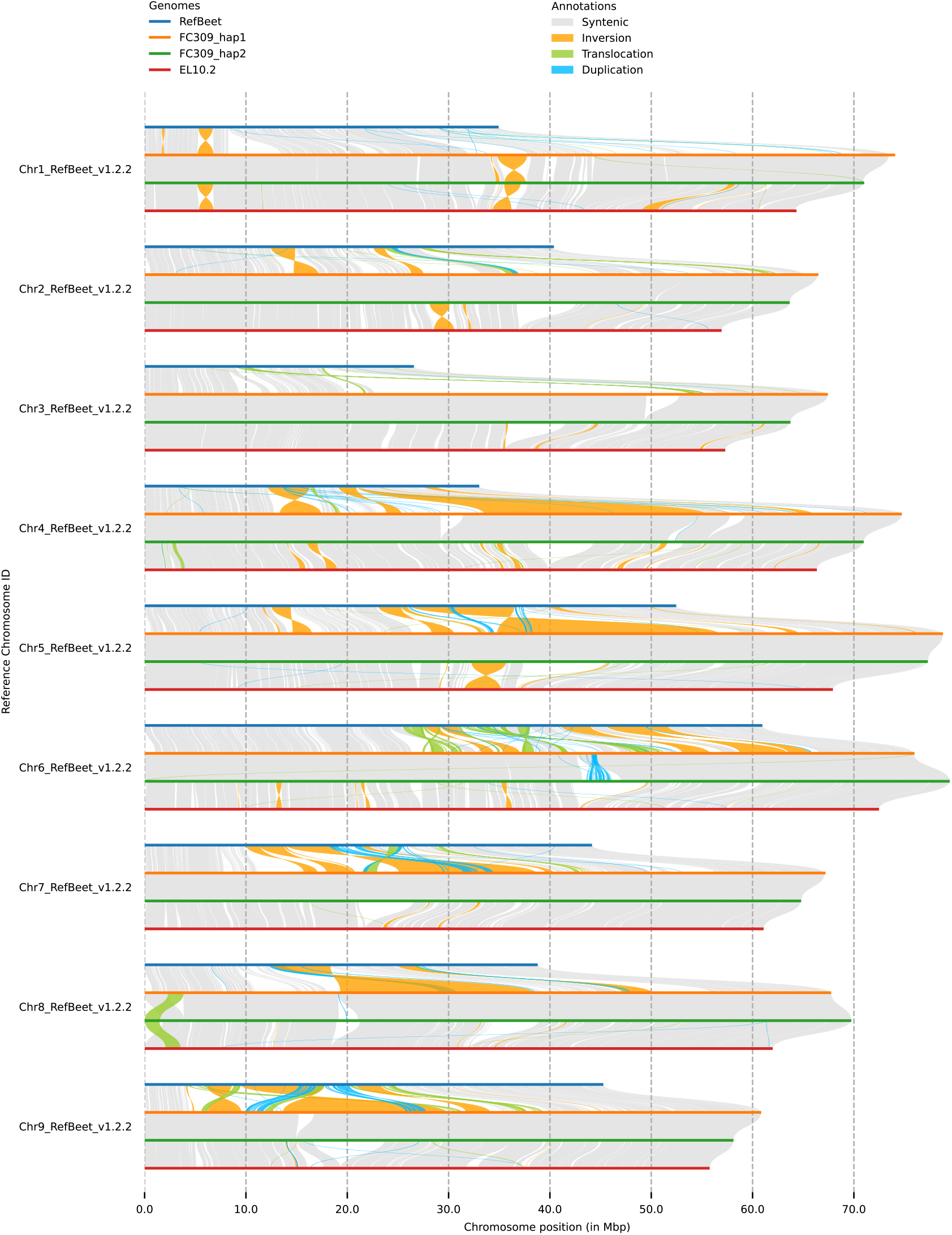
Genome-wide synteny analysis of FC309 phased assemblies, RefBeet (v1.2.2) and EL10.2. Chromosomes 1 through 9 are arranged vertically, with syntenic and variations between RefBeet (blue, top), FC309 v1.1.0 (orange), FC309 v1.2.0 (green), and EL10.2 (red, bottom) highlighted. Syntenic regions identified by Syri are represented by gray ribbons. Inversions (orange ribbons), transversions (green ribbons), and duplications (blue ribbons) are also highlighted for each chromosome comparison.

**Supplemental Figure 2:**
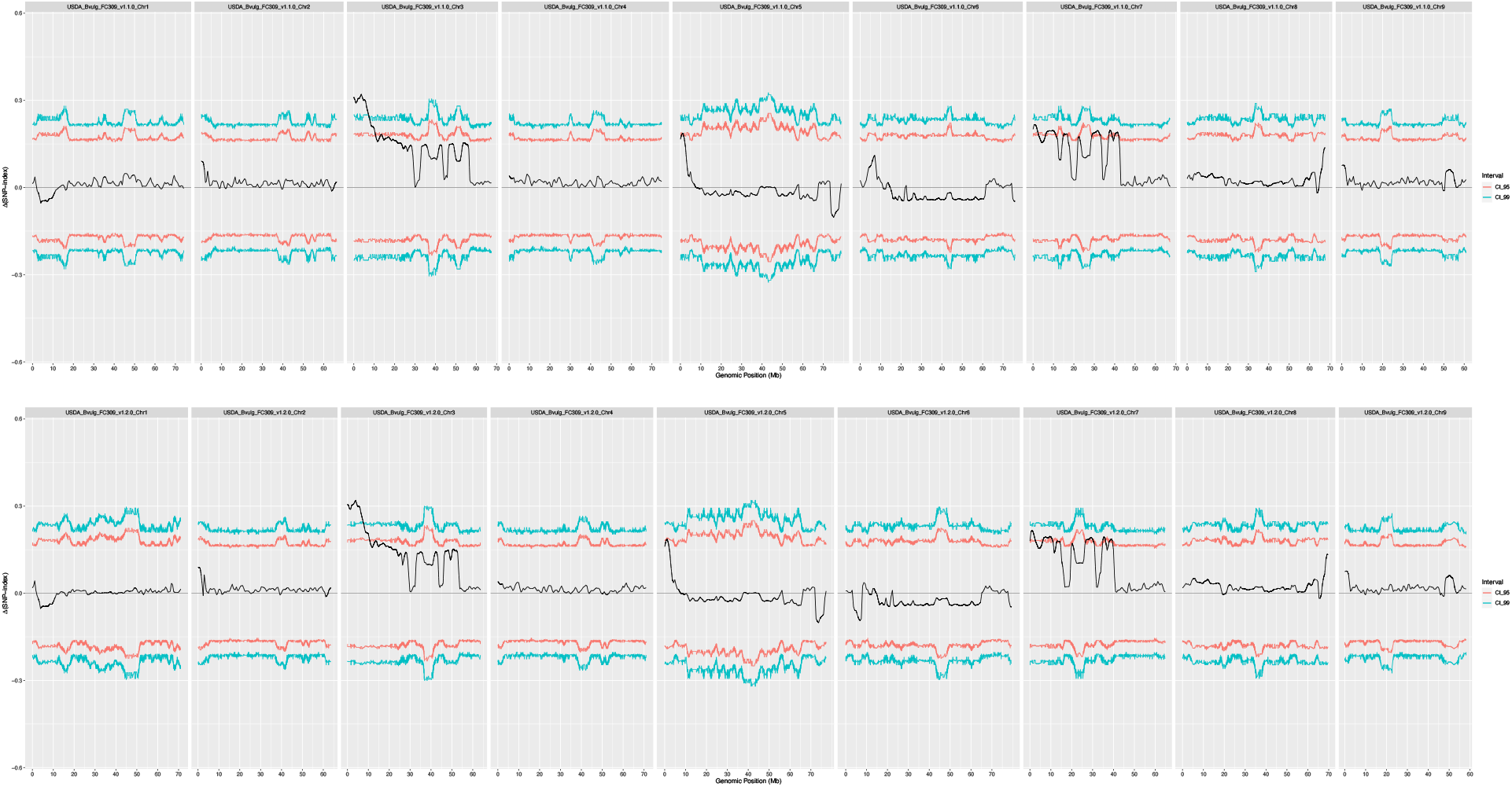
Tri-cubed ΔSNP index statistic (black line) of the *Fusarium* resistant vs. *Fusarium* susceptible FC308 x FC309 F2 mapping-by-sequencing pools mapped against the FC309 phased assemblies. Haplome 1 (v1.1.0) is shown on the top panel, haplome 2 (v1.2.0) is shown on the bottom panel, with each chromosome arranged left to right. 95% (red line) and 99% (blue line) confidence intervals are shown.

**Supplemental Table 1:** Repeat table generated by RepeatModeler.

| Class | Repeat Element | Number of elements |  | Length occupied (bp) |  | Percentage of sequence |  |
| --- | --- | --- | --- | --- | --- | --- | --- |
|  |  | Haplome 1 | Haplome 2 | Haplome 1 | Haplome 2 | Haplome 1 | Haplome 2 |
| **Retroelements** |  | **115917** | **127292** | **128988986** | **128489669** | **19.75** | **19.29** |
|  | SINEs | 1196 | 5529 | 2887779 | 4766107 | 0.44 | 0.72 |
|  | LINEs | 37705 | 25654 | 22453831 | 18616541 | 3.44 | 2.79 |
|  | RTE/Bov-B | 4374 | 932 | 1929389 | 335314 | 0.30 | 0.05 |
|  | L1/CIN4 | 33331 | 24722 | 20524442 | 18281227 | 3.14 | 2.74 |
|  | LTR elements | 77016 | 96109 | 103647376 | 105107021 | 15.87 | 15.78 |
|  | Ty1/Copia | 35144 | 34054 | 49418137 | 47739322 | 7.57 | 7.17 |
|  | Gypsy/DIRS1 | 40392 | 59726 | 52877555 | 55007311 | 8.10 | 8.26 |
|  | BEL/Pao | 0 | 229 | 0 | 24990 | 0 | 0.001 |
| **DNA transposons:** |  | **34421** | **24726** | **21245112** | **16970712** | **3.25** | **2.55** |
|  | hobo-Activator | 3647 | 4606 | 1822821 | 2447975 | 0.28 | 0.37 |
|  | Tc1-IS630-Pogo | 1174 | 1358 | 855393 | 923774 | 0.13 | 0.14 |
|  | Tourist/Harbinger | 3716 | 5280 | 1680204 | 2168591 | 0.26 | 0.33 |
| **Unclassified** |  | **710970** | **896950** | **261458489** | **256038339** | **40.04** | **38.44** |

**Supplemental Table 2:**
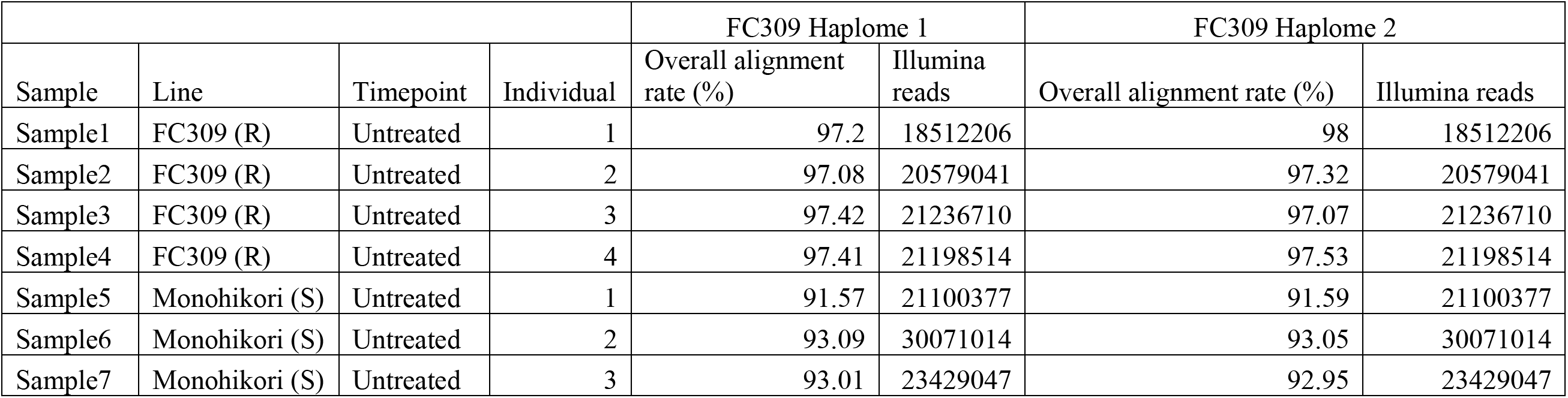
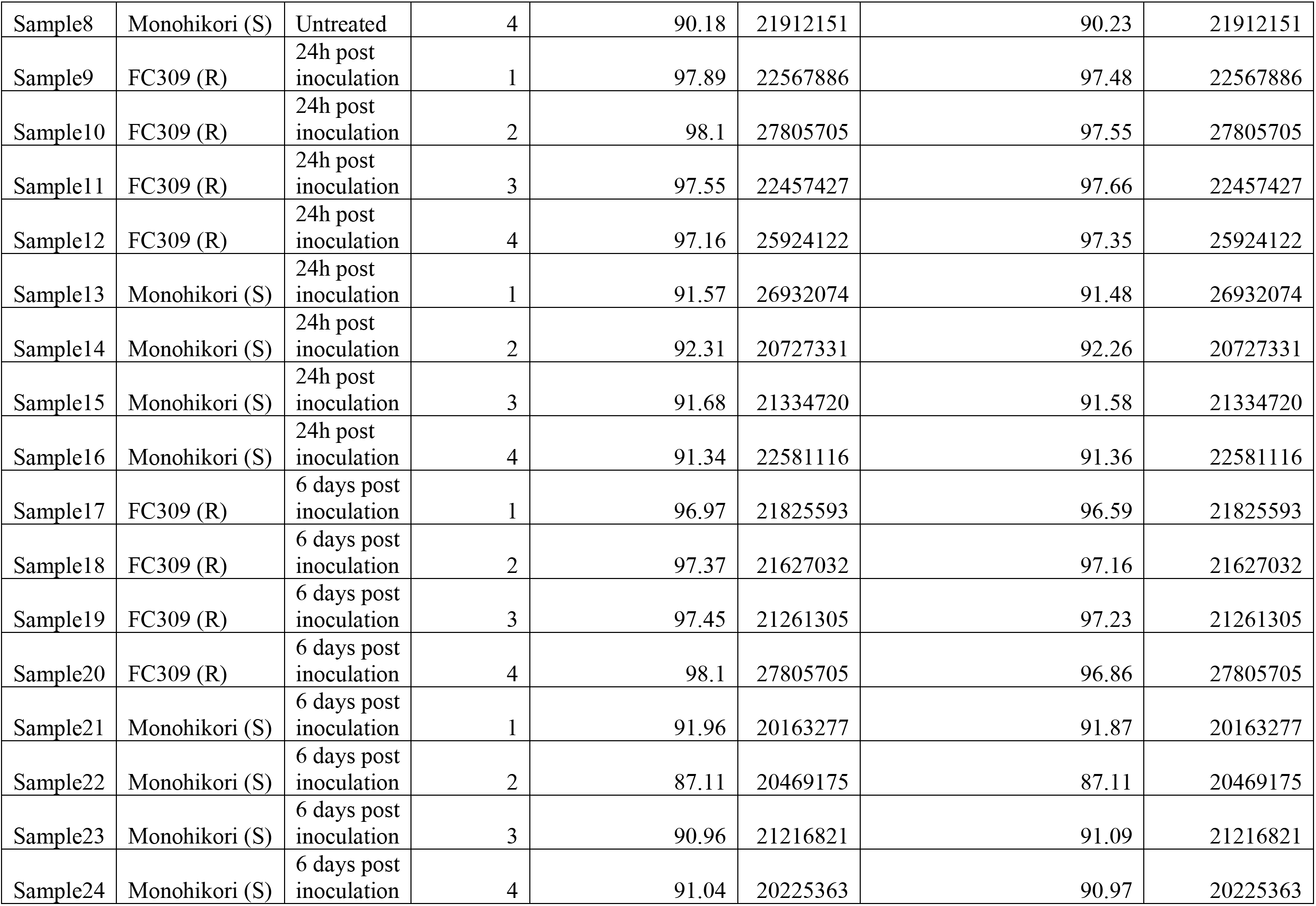
Sequencing and alignment statistics for the alignment of illumina reads to each assembled FC309 haplome for two lines, FC309 and Monohikori.

|  |  |  |  | FC309 Haplome 1 |  | FC309 Haplome 2 |  |
| --- | --- | --- | --- | --- | --- | --- | --- |
| Sample | Line | Timepoint | Individual | Overall alignment rate (%) | Illumina reads | Overall alignment rate (%) | Illumina reads |
| Sample1 | FC309 (R) | Untreated | 1 | 97.2 | 18512206 | 98 | 18512206 |
| Sample2 | FC309 (R) | Untreated | 2 | 97.08 | 20579041 | 97.32 | 20579041 |
| Sample3 | FC309 (R) | Untreated | 3 | 97.42 | 21236710 | 97.07 | 21236710 |
| Sample4 | FC309 (R) | Untreated | 4 | 97.41 | 21198514 | 97.53 | 21198514 |
| Sample5 | Monohikori (S) | Untreated | 1 | 91.57 | 21100377 | 91.59 | 21100377 |
| Sample6 | Monohikori (S) | Untreated | 2 | 93.09 | 30071014 | 93.05 | 30071014 |
| Sample7 | Monohikori (S) | Untreated | 3 | 93.01 | 23429047 | 92.95 | 23429047 |
| Sample8 | Monohikori (S) | Untreated | 4 | 90.18 | 21912151 | 90.23 | 21912151 |
| Sample9 | FC309 (R) | 24h post inoculation | 1 | 97.89 | 22567886 | 97.48 | 22567886 |
| Sample10 | FC309 (R) | 24h post inoculation | 2 | 98.1 | 27805705 | 97.55 | 27805705 |
| Sample11 | FC309 (R) | 24h post inoculation | 3 | 97.55 | 22457427 | 97.66 | 22457427 |
| Sample12 | FC309 (R) | 24h post inoculation | 4 | 97.16 | 25924122 | 97.35 | 25924122 |
| Sample13 | Monohikori (S) | 24h post inoculation | 1 | 91.57 | 26932074 | 91.48 | 26932074 |
| Sample14 | Monohikori (S) | 24h post inoculation | 2 | 92.31 | 20727331 | 92.26 | 20727331 |
| Sample15 | Monohikori (S) | 24h post inoculation | 3 | 91.68 | 21334720 | 91.58 | 21334720 |
| Sample16 | Monohikori (S) | 24h post inoculation | 4 | 91.34 | 22581116 | 91.36 | 22581116 |
| Sample17 | FC309 (R) | 6 days post inoculation | 1 | 96.97 | 21825593 | 96.59 | 21825593 |
| Sample18 | FC309 (R) | 6 days post inoculation | 2 | 97.37 | 21627032 | 97.16 | 21627032 |
| Sample19 | FC309 (R) | 6 days post inoculation | 3 | 97.45 | 21261305 | 97.23 | 21261305 |
| Sample20 | FC309 (R) | 6 days post inoculation | 4 | 98.1 | 27805705 | 96.86 | 27805705 |
| Sample21 | Monohikori (S) | 6 days post inoculation | 1 | 91.96 | 20163277 | 91.87 | 20163277 |
| Sample22 | Monohikori (S) | 6 days post inoculation | 2 | 87.11 | 20469175 | 87.11 | 20469175 |
| Sample23 | Monohikori (S) | 6 days post inoculation | 3 | 90.96 | 21216821 | 91.09 | 21216821 |
| Sample24 | Monohikori (S) | 6 days post inoculation | 4 | 91.04 | 20225363 | 90.97 | 20225363 |

**Supplemental Table 3:**
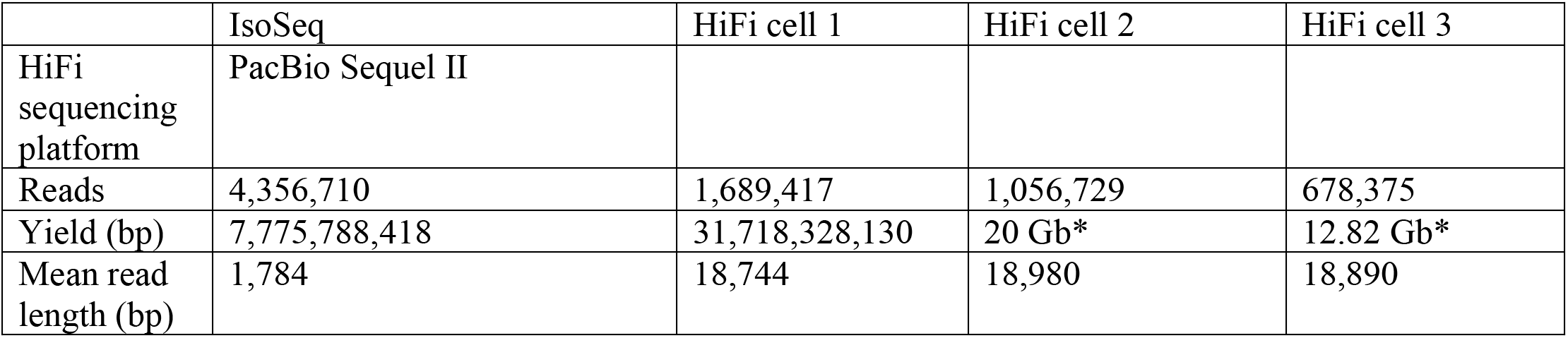
Isoseq and HiFi sequencing data statistics. *full sequencing report not available.

**Supplemental Table 4:**
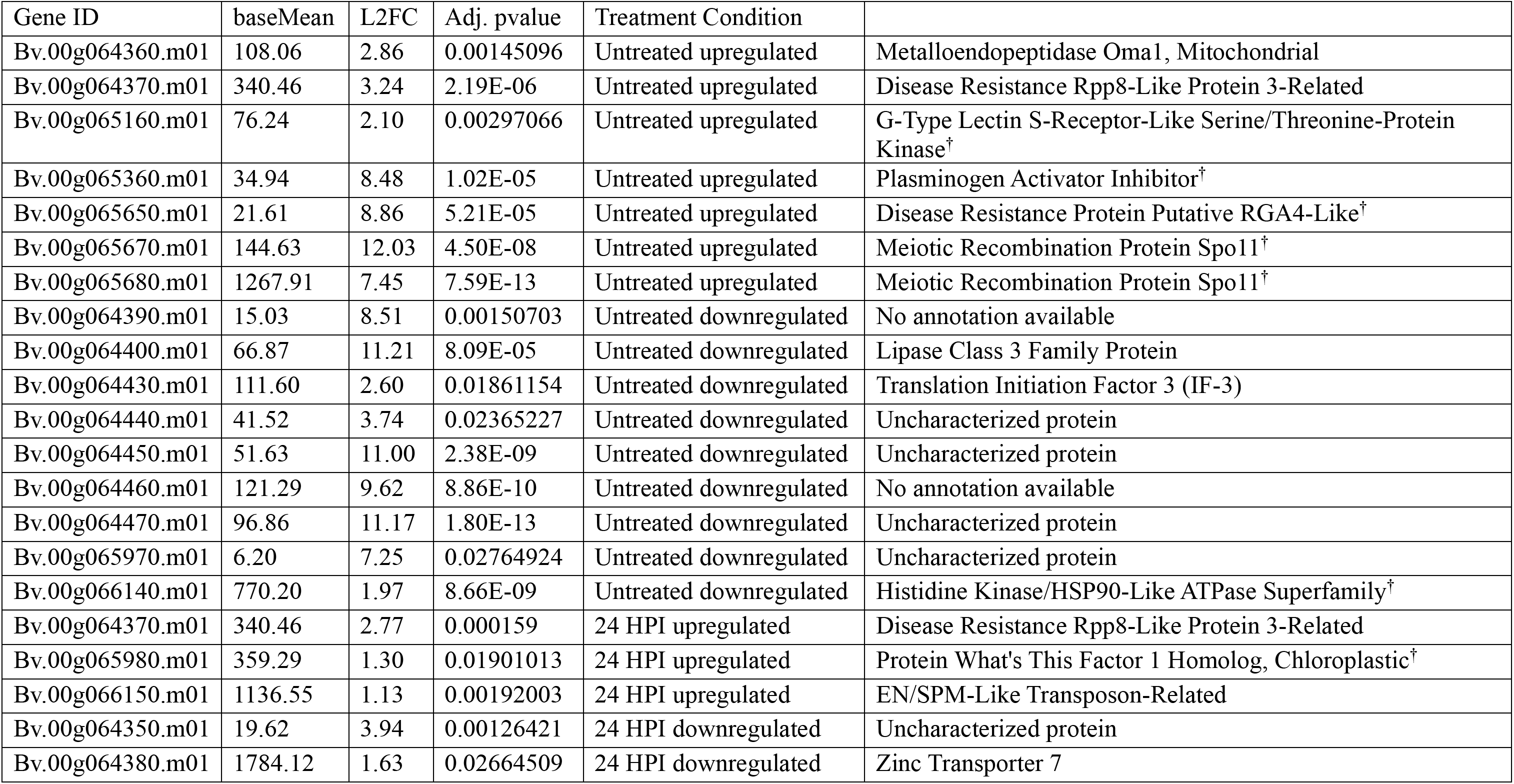
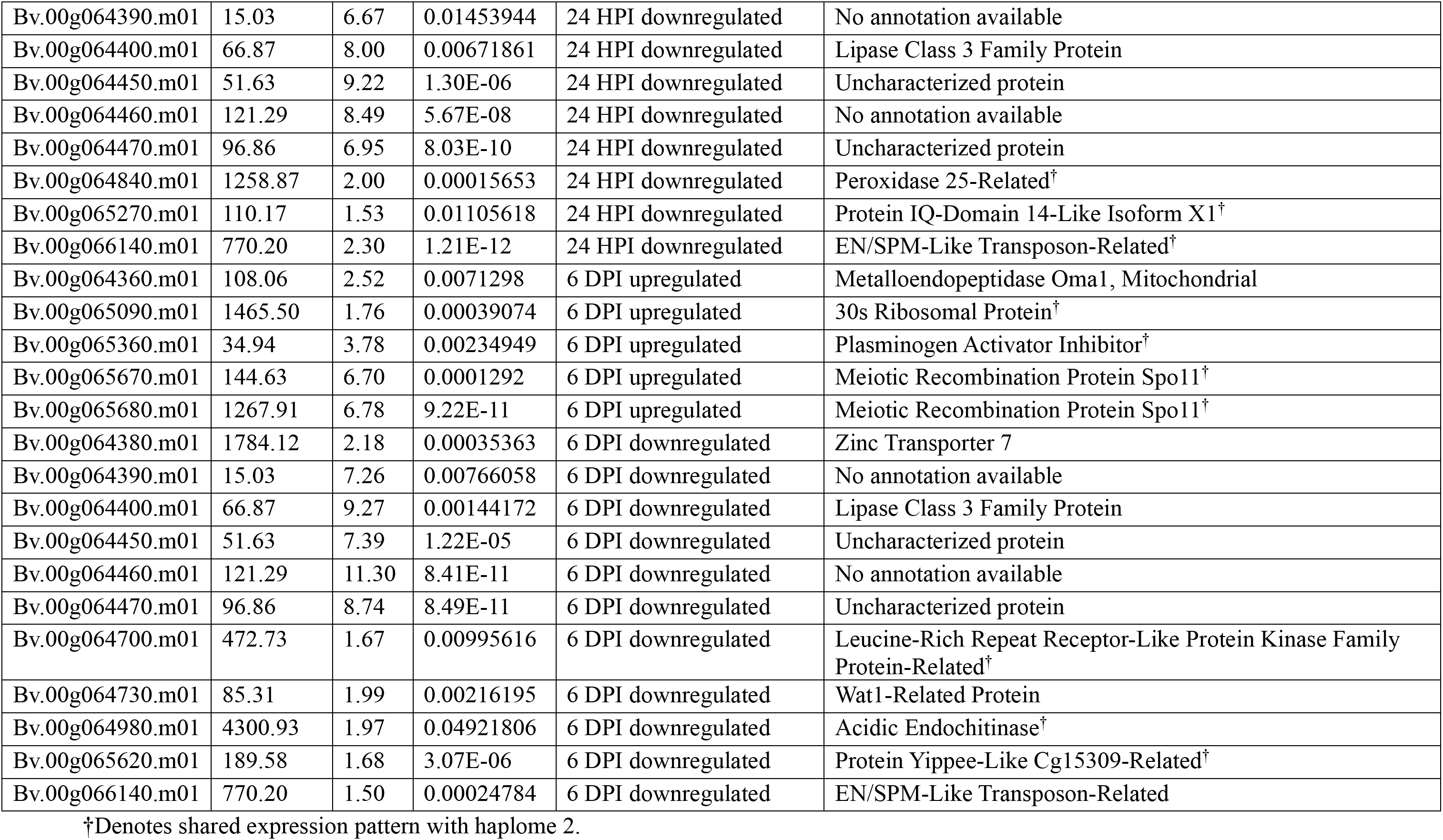
Haplome 1 differential transcript expression in the QTL region following RNA-sequencing. BaseMean represents the normalized count value over all samples output from DESeq2, log2 fold change (L2FC) is the absolute value of the change over the condition in the Treatment Condition column between the resistant and susceptible line (e.g. “Untreated upregulated” are the genes upregulated in the resistant line compared to the susceptible line at the untreated condition). Adj. Pvalues were adjusted using the Benjamini-Hochberg method to reduce false positives. InterPro annotations are provided by the InterPro consortium.

**Supplemental Table 5:** Haplome 2 differential transcript expression in the QTL region following RNA-sequencing. BaseMean represents the normalized count value over all samples output from DESeq2, log2 fold change (L2FC) is the absolute value of the change over the condition in the Treatment Condition column between the R and S line (e.g. “Untreated upregulated” are the genes upregulated in the resistant line compared to the susceptible line at the untreated condition). Pvalues were adjusted using the Benjamini-Hochberg method to reduce false positives. InterPro annotations are provided by the InterPro consortium.

| Gene ID | baseMean | L2FC | Adj. Pvalue | Treatment Condition | Annotation |
| --- | --- | --- | --- | --- | --- |
| Bv.00g064490.m01 | 27.49 | 6.34 | 0.0043977 | Untreated upregulated | Heavy Metal Transport/Detoxification Superfamily Protein |
| Bv.00g064870.m01 | 76.18 | 2.10 | 0.00305379 | Untreated upregulated | G-Type Lectin S-Receptor-Like Serine/Threonine-Protein Kinase<sup>†</sup> |
| Bv.00g064920.m01 | 9.63 | 6.56 | 0.00093816 | Untreated upregulated | Plastid Movement Impaired Protein-Related |
| Bv.00g065060.m01 | 34.98 | 8.49 | 7.54E-06 | Untreated upregulated | Plasminogen Activator Inhibitor<sup>†</sup> |
| Bv.00g065330.m01 | 21.57 | 8.86 | 4.98E-05 | Untreated upregulated | Disease Resistance Protein Putative RGA4-Like<sup>†</sup> |
| Bv.00g065360.m01 | 137.42 | 11.96 | 1.80E-08 | Untreated upregulated | Meiotic Recombination Protein Spo11<sup>†</sup> |
| Bv.00g065370.m01 | 1266.94 | 7.45 | 9.70E-13 | Untreated upregulated | Meiotic Recombination Protein Spo11<sup>†</sup> |
| Bv.00g066060.m01 | 33.80 | 2.10 | 0.03491854 | Untreated upregulated | Galactose Oxidase/Kelch, Beta-Propeller Superfamily |
| Bv.00g064410.m01 | 10.53 | 7.46 | 0.01332745 | Untreated downregulated | Reverse Transcriptase Zinc-Binding Domain-Containing Protein-Related |
| Bv.00g065660.m01 | 10.61 | 8.59 | 0.0036312 | Untreated downregulated | Modifying Wall Lignin-1/2 |
| Bv.00g065840.m01 | 786.36 | 2.00 | 6.25E-09 | Untreated downregulated | Histidine Kinase/HSP90-Like ATPase Superfamily<sup>†</sup> |
| Bv.00g064490.m01 | 27.49 | 2.78 | 0.04946334 | 24 HPI upregulated | Copper Transport Protein Atox1-Related |
| Bv.00g064920.m01 | 9.63 | 3.21 | 0.02022821 | 24 HPI upregulated | Movement Impaired Protein-Related |
| Bv.00g065060.m01 | 34.98 | 2.48 | 0.04949785 | 24 HPI upregulated | Plasminogen Activator Inhibitor |
| Bv.00g065680.m01 | 365.23 | 1.33 | 0.01258339 | 24 HPI upregulated | Protein What's This Factor 1 Homolog, Chloroplastic<sup>†</sup> |
| Bv.00g064550.m01 | 1259.12 | 2.00 | 0.0001443 | 24 HPI downregulated | Peroxidase 25-Related<sup>†</sup> |
| Bv.00g064970.m01 | 110.47 | 1.52 | 0.01025766 | 24 HPI downregulated | Protein IQ-Domain 14-Like Isoform X1<sup>†</sup> |
| Bv.00g065840.m01 | 786.36 | 2.39 | 1.32E-13 | 24 HPI downregulated | EN/SPM-Like Transposon-Related<sup>†</sup> |
| Bv.00g065950.m01 | 132.39 | 4.41 | 0.00049988 | 24 HPI downregulated | Alpha/Beta-Hydrolases Superfamily Protein |
| Bv.00g067220.m01 | 27.10 | 9.40 | 2.76E-07 | 24 HPI downregulated | Exosome Complex Exonuclease Ribosomal RNA Processing Protein |
| Bv.00g067690.m01 | 22.96 | 7.71 | 0.00218019 | 24 HPI downregulated | 3-Oxo-Delta(4,5)-Steroid 5-Beta-Reductase-Like |
| Bv.00g064490.m01 | 27.49 | 3.65 | 0.00834525 | 6 DPI upregulated | Copper Transport Protein Atox1-Related |
| Bv.00g064810.m01 | 1463.55 | 1.76 | 0.00038868 | 6 DPI upregulated | 30s Ribosomal Protein<sup>†</sup> |
| Bv.00g065060.m01 | 34.98 | 3.70 | 0.00232944 | 6 DPI upregulated | Plasminogen Activator Inhibitor<sup>†</sup> |
| Bv.00g065360.m01 | 137.42 | 10.98 | 4.06E-07 | 6 DPI upregulated | Meiotic Recombination Protein Spo11<sup>†</sup> |
| Bv.00g065370.m01 | 1266.94 | 6.77 | 1.18E-10 | 6 DPI upregulated | Meiotic Recombination Protein Spo11<sup>†</sup> |
| Bv.00g064380.m01 | 473.15 | 1.67 | 0.01014084 | 6 DPI downregulated | Leucine-Rich Repeat Receptor-Like Protein Kinase Family Protein-Related<sup>†</sup> |
| Bv.00g064410.m01 | 10.53 | 7.48 | 0.01070472 | 6 DPI downregulated | Reverse Transcriptase Zinc-Binding Domain-Containing Protein-Related |
| Bv.00g064710.m01 | 4302.55 | 1.98 | 0.04733243 | 6 DPI downregulated | Acidic Endochitinase<sup>†</sup> |
| Bv.00g065300.m01 | 190.35 | 1.73 | 5.57E-06 | 6 DPI downregulated | Protein Yippee-Like Cg15309-Related<sup>†</sup> |
| Bv.00g065840.m01 | 786.36 | 1.53 | 0.0001772 | 6 DPI downregulated | ATPase Domain Of HSP90 Chaperone/DNA Topoisomerase II/Histidine Kinase |
| Bv.00g066040.m01 | 3444.87 | 1.01 | 4.09E-05 | 6 DPI downregulated | Zinc Finger Protein Constans-Like 10 |

<sup>†</sup>Denotes shared expression pattern with haplome 1.

**Table 6:** Combined transcript expression comparisons in the QTL region following RNA-sequencing within the resistant line over both the 24 hours post inoculaton (HPI) and 6 days post inoculation (DPI) compared to the untreated timepoint. BaseMean represents the normalized count value over all samples output from DESeq2, log2 fold change (L2FC) is presented as the absolute value of the change over the condition in the Treatment Condition within the resistant line (e.g. “24 HPI upregulated” are the genes upregulated at the 24 HPI timepoint compared to the untreated timepoint in resistant line). Pvalues were adjusted using the Benjamini-Hochberg method to reduce false positives. InterPro annotations are provided by the InterPro consortium. Gene IDs are from the haplome 2 annotation unless otherwise denoted. All transcripts were similarly expressed between haplome 1 and 2 unless otherwise denoted.

| Gene ID | baseMean | L2FC | Adj. Pvalue | Treatment Condition | Annotation |
| --- | --- | --- | --- | --- | --- |
| Bv.00g064370.m01 | 718.93 | 2.42 | 1.27E-09 | 24 HPI upregulated | L-Type Lectin-Domain Containing Receptor Kinase VIII.2-Like |
| Bv.00g064680.m01 | 10039.80 | 1.05 | 0.001508773 | 24 HPI upregulated | Bzip Transcription Factor Bzip124<sup>2</sup> |
| Bv.00g064960.m01 | 638.77 | 6.44 | 2.00E-26 | 24 HPI upregulated | Heat Shock Protein 21, Chloroplastic |
| Bv.00g065070.m01 | 2418.53 | 1.41 | 0.009631139 | 24 HPI upregulated | Homeobox-Leucine Zipper Protein Athb-7 |
| Bv.00g065190.m01 | 151.08 | 2.76 | 2.53E-06 | 24 HPI upregulated | Centromere Protein V<sup>2</sup> |
| Bv.00g065400.m01 | 553.13 | 1.18 | 0.000663691 | 24 HPI upregulated | F-Box Only Protein 48-Related |
| Bv.00g065890.m01 | 5338.59 | 0.91 | 0.01144541 | 24 HPI upregulated | Clp Protease Regulatory Subunit Clpx1, Mitochondrial |
| Bv.00g065510.m01 | 123.33 | 2.69 | 5.61E-06 | 24 HPI upregulated | Uncharacterized protein<sup>1</sup> |
| Bv.00g064190.m01 | 684.21 | 1.06 | 0.001571901 | 24 HPI upregulated | Modifying Wall Lignin-1/2<sup>1</sup> |
| Bv.00g064550.m01 | 1259.12 | 2.77 | 5.99E-10 | 24 HPI downregulated | Peroxidase 25-Related |
| Bv.00g064560.m01 | 697.45 | 1.90 | 0.002313416 | 24 HPI downregulated | Ankyrin Repeat Protein-Like |
| Bv.00g064760.m01 | 998.63 | 1.44 | 0.001964188 | 24 HPI downregulated | Syntaxin |
| Bv.00g064850.m01 | 442.11 | 3.24 | 1.45E-11 | 24 HPI downregulated | Avr9/Cf-9 Rapidly Elicited Protein |
| Bv.00g064980.m01 | 792.24 | 2.37 | 2.63E-13 | 24 HPI downregulated | Growth-Regulating Factor 7 |
| Bv.00g065260.m01 | 1296.10 | 3.72 | 1.13E-20 | 24 HPI downregulated | Oligopeptide Transporter 6 |
| Bv.00g065650.m01 | 6689.80 | 3.07 | 1.67E-15 | 24 HPI downregulated | Modifying Wall Lignin-1/2 |
| Bv.00g065690.m01 | 106.61 | 2.94 | 0.006236752 | 24 HPI downregulated | Modifying Wall Lignin-1/2 |
| Bv.00g065700.m01 | 2242.45 | 1.50 | 8.01E-07 | 24 HPI downregulated | Modifying Wall Lignin-1/2 |
| Bv.00g065900.m01 | 1578.51 | 1.79 | 1.13E-06 | 24 HPI downregulated | Calcineurin B-Like Protein 4 |
| Bv.00g065940.m01 | 1436.00 | 1.59 | 0.002524712 | 24 HPI downregulated | Pectinesterase/Pectinesterase Inhibitor 34-Related<sup>2</sup> |
| Bv.00g064330.m01 | 60.05 | 4.45 | 0.000831819 | 6 DPI upregulated | Metalloendopeptidase Oma1<sup>1</sup> |
| Bv.00g064250.m01 | 3092.16 | 0.86 | 0.041423882 | 6 DPI upregulated | Expressed Protein<sup>2</sup> |
| Bv.00g064420.m01 | 25.95 | 1.89 | 0.045631506 | 6 DPI upregulated | Transcription Repressor Ofp7-Related<sup>2</sup> |
| Bv.00g064810.m01 | 1463.55 | 2.24 | 3.62E-07 | 6 DPI upregulated | 30s Ribosomal Protein<sup>2</sup> |
| Bv.00g064850.m01 | 442.11 | 2.06 | 0.000409659 | 6 DPI downregulated | Avr9/Cf-9 Rapidly Elicited Protein |
| Bv.00g065540.m01 | 17215.83 | 0.86 | 0.031445793 | 6 DPI downregulated | Heat Shock Protein 70kd |
| Bv.00g065750.m01 | 1633.65 | 1.44 | 0.009534603 | 6 DPI downregulated | Ferrochelatase-1, Chloroplastic/Mitochondrial |
| Bv.00g066040.m01 | 3444.87 | 1.13 | 1.22E-07 | 6 DPI downregulated | Zinc Finger Protein Constans-Like 10<sup>2</sup> |

<sup>1</sup>Denotes presence in QTL region of haplome 1 only
<sup>2</sup>Denotes presence in QTL region of haplome 2 only

